# Cyclosporin A delays the terminal disease stage in *Tfam* KO mice without improving mitochondrial energy production

**DOI:** 10.1101/2022.10.14.511701

**Authors:** Benjamin Chatel, Isabelle Varlet, Augustin C. Ogier, Emilie Pecchi, Monique Bernard, Julien Gondin, Håkan Westerblad, David Bendahan, Charlotte Gineste

**Author notes:** **Corresponding Author:** Charlotte GINESTE; IGBMC, 1 Rue Laurent Fries, 67400 Illkirch, France; 00 33 (0)3 88 65 34 14.

## Abstract

Mitochondrial myopathies are rare genetic disorders characterized by muscle weakness and exercise intolerance. Currently, no effective treatment exists for these myopathies. Interestingly, the pharmacological cyclophilin inhibitor cyclosporine A (CsA) extended lifespan and prevented loss of force and mitochondrial Ca^2+^ overload in muscle fibers in the skeletal muscle-specific *Tfam* knockout mouse model of lethal mitochondrial myopathy (*Tfam* KO). The unaffected expression of proteins involved in mitochondrial energy metabolism suggests that these improvements occurred without improvement in metabolism. In this study, we aimed at investigating the effects of four weeks of CsA administration on *in vivo* contractile function and mitochondrial energy production in *Tfam* KO mice. The treatment started before the terminal phase with severe muscle weakness and weight loss. Our results show that CsA treatment delayed progression into the terminal disease phase. This occurred without any obvious positive effects on mitochondrial energy production at rest or during fatigue induced by repeated contractions. In conclusion, cyclophilin inhibitors may have the potential of counteracting devastating muscle weakness in patients with mitochondrial myopathies most probably by preventing deleterious effects triggered by excessive mitochondrial Ca^2+^ uptake rather than by improving mitochondrial energy production.

## Introduction

Mitochondrial myopathies compose a heterogeneous group of genetic diseases often resulting in the failure to generate the required cellular energy by oxidative phosphorylation (1). Patients with mitochondrial disease present a range of symptoms and clinical manifestations including muscle wasting, muscle weakness and exercise intolerance (2–4). No effective therapy is available and agents aiming at increasing respiratory chain components, ketogenic diet, amino-acids supplementation and exercise are the main therapeutic options used to alleviate the muscle symptoms (5–8). Furthermore, the complex and variable phenotypes of mitochondrial myopathies make it difficult to assess the efficacy of a treatment (9, 10).

Mice with skeletal muscle-specific deletion of the gene for mitochondrial transcription factor A (*Tfam* KO mice) faithfully reproduce the main structural and clinical features of mitochondrial myopathy, *i. e*. muscle atrophy, ragged red fibers, muscle weakness, muscle fatigue and mitochondrial dysfunction (11, 12). An abnormally large and slowly reversible increase in mitochondrial Ca^2+^ was observed during repeated contractions in *Tfam* KO muscle fibers and this was suggested to play a main role in the disease process (13). This excessive mitochondrial Ca^2+^ uptake was partially inhibited by cyclosporin A (CsA) (13). CsA is a pharmacological agent that binds to the mitochondrial matrix protein peptidyl-prolyl cis-trans isomerase F (aka cyclophilin D, CypD) and is considered to desensitize opening of the mitochondrial permeability transition pore (14). In *Tfam* KO mice, CsA treatment counteracted the rapidly progressing terminal muscle weakness and extended the lifespan (15). Interestingly, the protein expression of CypD has been shown to be increased in *Tfam* KO muscles and also in patients with mitochondrial myopathies (15), thus implying that treatment with CsA might be effective in combating muscle defects in this pathology.

We recently showed that impaired mitochondrial energy production and premature fatigue occur before *Tfam* KO mice enter the final terminal state with rapidly developing muscle weakness, weight loss and ultimately early death (11). These results agree with previous data obtained in mouse models with mitochondrial dysfunction, as well as in patients with mitochondrial myopathies (16–18). Intriguingly, the CsA treatment-induced protection against muscle weakness, weight loss and early death in *Tfam* KO mice was not accompanied by any obvious signs of improved mitochondrial respiration, as judged from unaffected expression of proteins involved in mitochondrial energy metabolism (15). This intricate finding implies that mitochondrial myopathies might be effectively treated with pharmacological agents that do not improve mitochondrial respiration. Nevertheless, this counterintuitive implication clearly requires further scientific support.

Thus, we here hypothesized that CsA treatment of *Tfam* KO mice could delay the disease progression into the terminal state without any beneficial effects on *in vivo* muscle fatigue and mitochondrial energy production. CsA treatment started when mice were 12-13 weeks old (*i. e*. before the *Tfam* KO mice displayed any obvious muscle weakness or weight loss) and lasted for 4 weeks. In accordance with our hypothesis, the results show delayed weight loss with CsA treatment, whereas no improvement in parameters related to muscle fatigue development and mitochondrial energy production was observed.

## Results

### CsA treatment did not affect contractile function or metabolism in WT mice

Four weeks of CsA treatment of WT mice had no significant effect on muscle volume, maximal tetanic force in the unfatigued state or the force produced during fatiguing stimulation (Supplementary Figure 1). Likewise, CsA treatment did not significantly affect ^31^P-MRS measurements of metabolites either in the rested state (Supplementary Figure 2) or during induction of fatigue and the subsequent recovery period (Supplementary Figure 3) in WT mice. Thus, pooled mean data from CsA-treated and untreated WT mice were used for comparison with results obtained in *Tfam* KO mice.

### CsA treatment delayed the decline in body weight in Tfam KO mice

*Tfam* KO mice rapidly lose weight when they enter the terminal state of myopathy (12), and CsA treatment has been shown to spare muscle mass and increase the lifespan of *Tfam* KO mice (15). The body weight at the start of the intervention period was similar for CsA-treated (20.7±0.7 g) and untreated (21.9±0.8 g) *Tfam* KO mice. Body weight was measured every 2-3 days from the start of intervention and CsA-treated *Tfam* KO mice reached their maximal body weight about 3 weeks later than untreated mice (Figure 1A). The untreated *Tfam* KO mice started to lose weight after the intervention period was initiated. In order to test untreated *Tfam* KO mice before endpoint was reached (*i. e*. >15% body weight loss), experiments were performed at 14 weeks in this group of mice. The CsA-treated *Tfam* KO mice were tested at the end of the 4-week administration period, *i. e*. at 16 weeks. Therefore, CsA-treated *Tfam* KO mice were older at the time of anatomical and functional testing as compared to untreated *Tfam* KO mice (Figure 1B). Despite CsA-treated mice being older at the time of testing, their body weight (expressed as per cent of that at the start of the intervention) was significantly higher than for untreated mice (Figure 1C). Thus, the 4 weeks of CsA treatment delayed the progressive decline in body weight, which is a key feature when *Tfam* KO mice are about to enter the terminal myopathy state.

**Figure 1.**
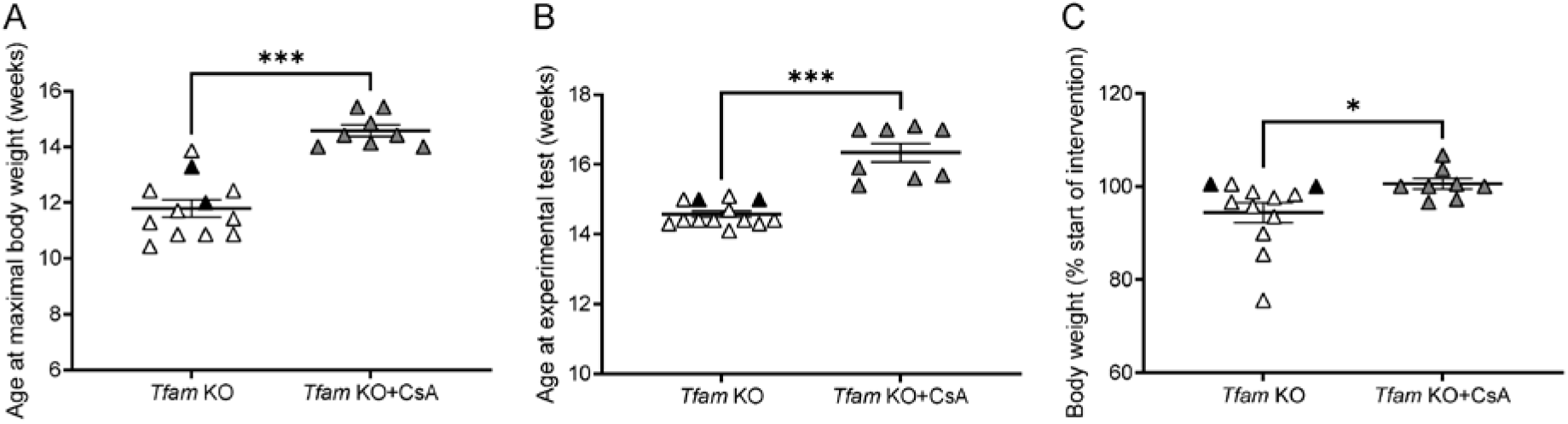
The time at which *Tfam* KO mice enters severe stage is delayed following CsA administration. **(A)** Age at which maximal body weight was reached. **(B)** Age when experimental procedures where applied. **(C)** Change in body mass from the intervention onset. Data presented as individual values and mean±SEM. *Tfam* KO, n=12-13; *Tfam* KO+CsA, n=11-12. White=mouse without pump; Black=mouse with pump+placebo; Grey=mouse with pump+CsA. Significant difference * and *** *P*<0.05 and 0.001, respectively; unpaired t-test.

MRI scanning showed that hindlimb muscles volume was similarly decreased in CsA-treated and untreated *Tfam* KO mice as compared to WT mice (~15%; Figure 2, A and B). The specific maximal tetanic (150 Hz) force in the unfatigued state was similar in WT, CsA-treated and untreated *Tfam* KO mice (Figure 2C).

**Figure 2.**
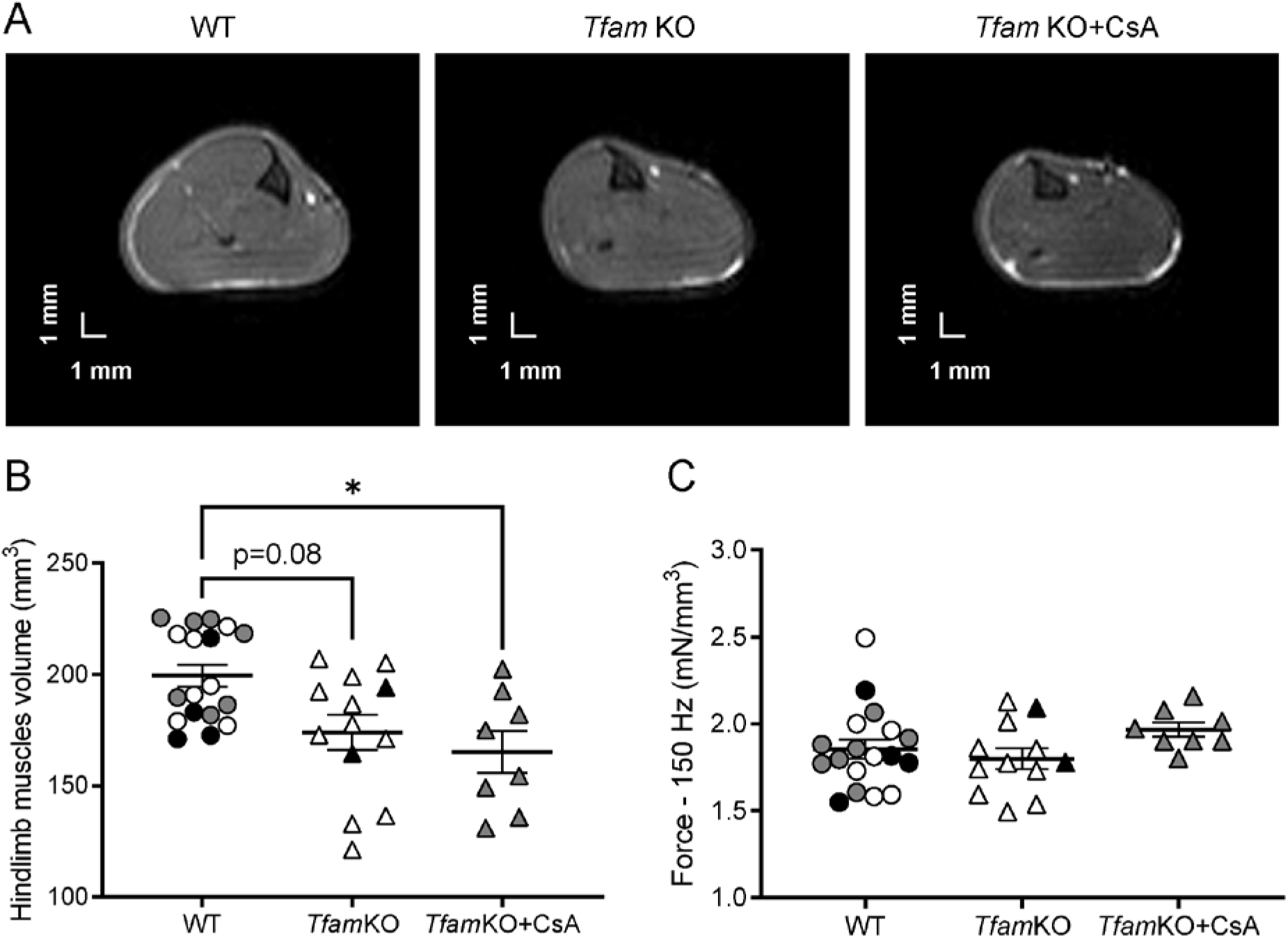
CsA administration did not affect muscle atrophy and maximal force in *Tfam* KO mice. Representative axial magnetic resonance images **(A)** and quantification of the volume of hindlimb muscles **(B). (C)** Maximal specific force of the plantar flexor muscles *in vivo*. Data presented as individual values and mean±SEM. WT, n=19; *Tfam* KO, n=12-13; *Tfam* KO+CsA, n=11. White=mouse without pump; Black=mouse with pump+placebo; Grey=mouse with pump+CsA. Significant difference * *P*<0.05; Kruskal-Wallis test and Dunn’s post hoc test for panel B; one-way ANOVA and Tuckey’s post hoc test for panel C.

### Neither the faster fatigue development nor the metabolic impairments in Tfam KO mice were prevented by CsA administration

We recently demonstrated that during fatiguing stimulation, skeletal muscle force declined rapidly due to impaired mitochondrial energy metabolism in *Tfam* KO mice (11). In the present study we assessed whether administration of CsA would counteract these deficiencies. The decline in force production during fatiguing stimulation was similar in CsA-treated and untreated *Tfam* KO mice, and markedly faster than in WT mice (Figure 3A). Accordingly, the overall force produced during the stimulation protocol and the C_50_ (number of contractions at which force was 50% of the initial) were not different in CsA-treated and untreated *Tfam* KO mice and significantly lower than in WT mice (Figure 3, B and C). Overall, no positive effect of CsA administration could be disclosed for muscle fatigue in *Tfam* KO mice.

**Figure 3.**
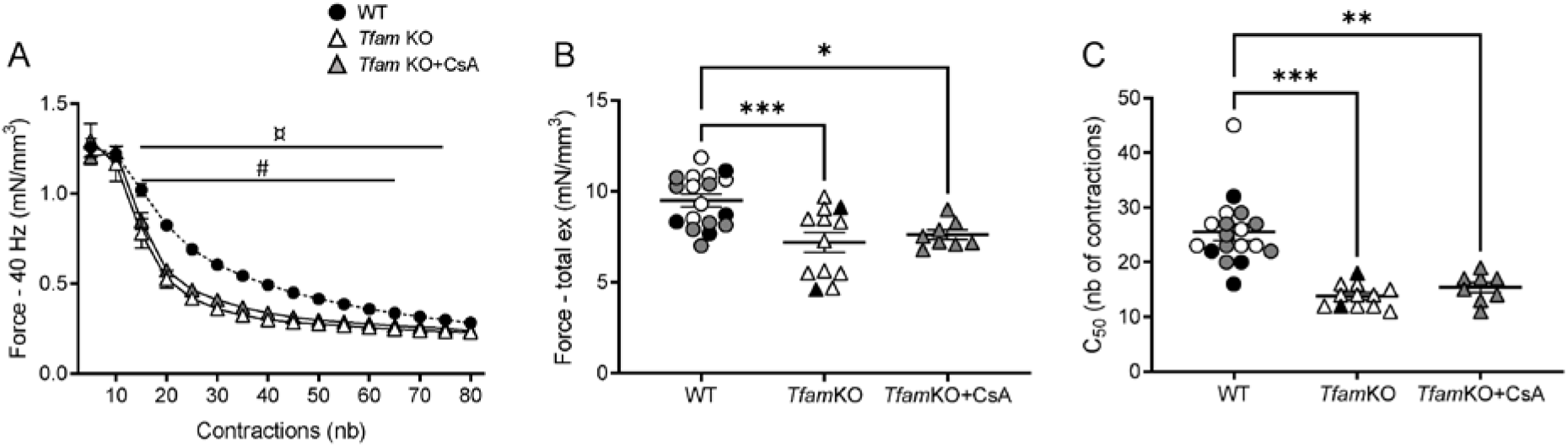
CsA had no effect on muscle fatigue in *Tfam* KO mice. **(A)** *In vivo* specific force of the plantar flexor muscles during fatiguing exercise. **(B)** Total specific force of the plantar flexor muscles produced during the whole fatiguing protocol. **(C)** Contraction at which force production was 50% of the unfatigued force (C_50_). Data presented as mean±SEM for panel A. Data presented as individual values and mean±SEM for panels B and C. WT, n=19; *Tfam* KO, n=12; *Tfam* KO+CsA, n=11. White=mouse without pump; Black=mouse with pump+placebo; Grey=mouse with pump+CsA for panels B and C. Significant difference between WT and *Tfam* KO ^¤^ *P*<0.05. Significant difference between WT and *Tfam* KO+CsA ^#^ *P*<0.05. Significant difference *, ** and *** *P*<0.05, 0.01 and 0.001, respectively. Two-way ANOVA with repeated measures on contraction number and Sidak’s post hoc test for panel A; one-way ANOVA and Tuckey’s post hoc test for panel B; Kruskal-Wallis test and Dunn’s post hoc test for panel C.

At rest, ^31^P-MRS measurements showed lower phosphocreatine (PCr) to ATP ratio (although not statistically significant in the untreated *Tfam* KO group), higher concentration of inorganic phosphate (Pi; expressed as the Pi/(Pi + PCr) ratio), and decreased pH_i_ in *Tfam* KO than in WT mice (Figure 4). All these measurements indicate a defective mitochondrial energy production in *Tfam* KO mice already at rest and CsA treatment did not reverse this deficiency.

**Figure 4.**
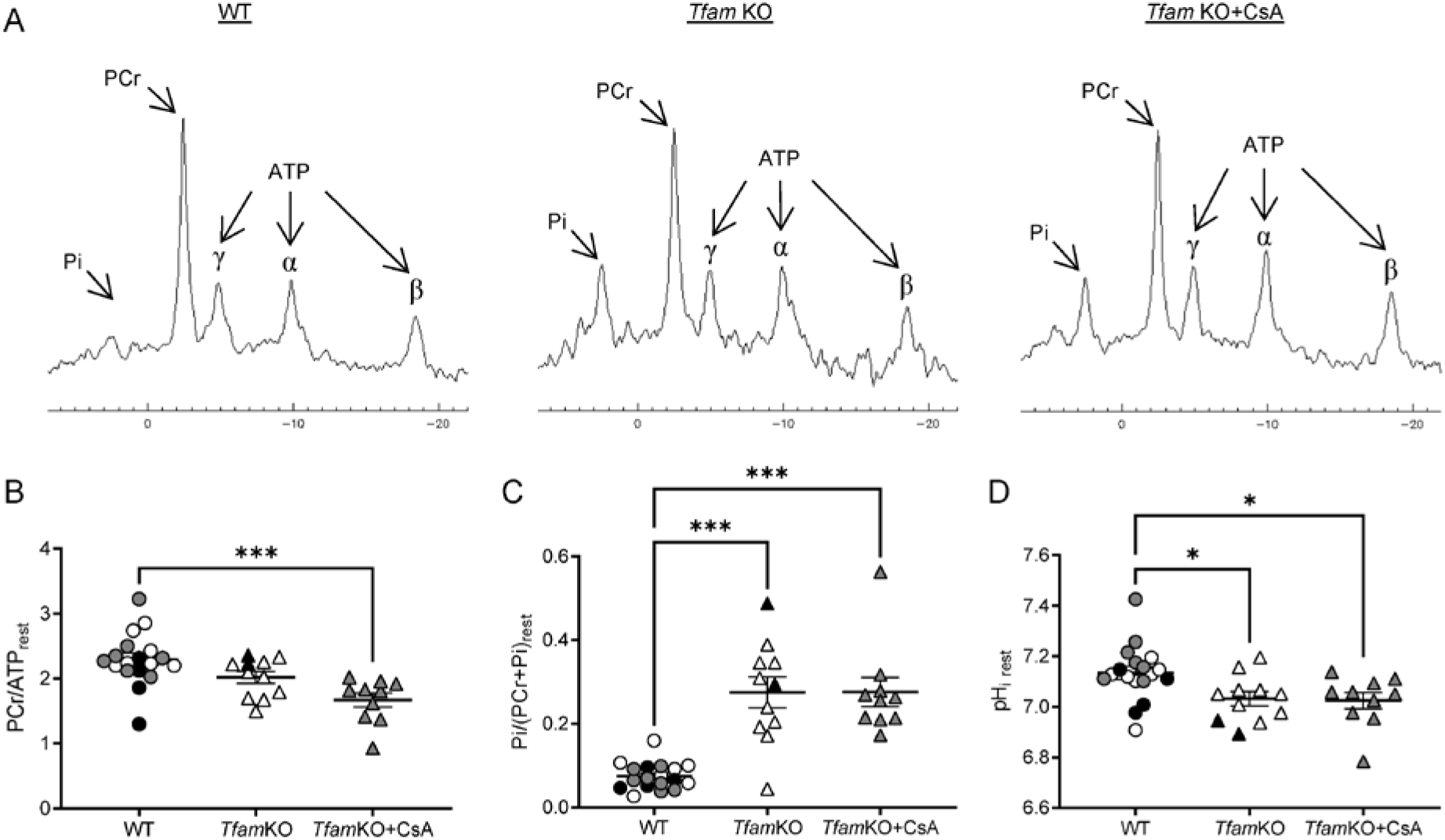
Metabolic parameters at rest were unchanged in *Tfam* KO mice after 4-week administration of CsA. **(A)** Example of resting ^31^P-MR spectra in WT (left panel), untreated *Tfam* KO (middle panel) and CsA-treated *Tfam* KO (right panel) muscles. Phosphocreatine to ATP ratio (**B**), inorganic phosphate (**C**) and pH_i_ (**D**) measured at rest. Data presented as individual values and mean±SEM. WT, n=19; *Tfam* KO, n=11; *Tfam* KO+CsA, n=10. White=mouse without pump; Black=mouse with pump+placebo; Grey=mouse with pump+CsA. Significant difference * and *** *P*<0.05 and 0.001, respectively. One-way ANOVA and Tuckey’s post hoc test for panels B and D; Kruskal-Wallis test and Dunn’s post hoc test for panel C. PCr: phosphocreatine; ATP: Adenosine tri-phosphate; Pi: inorganic phosphate; pH_i_: intracellular pH.

During fatiguing stimulation, the extent of PCr depletion was larger and Pi reached a higher level in *Tfam* KO than in WT mice (Figure 5, A-D). On the other hand, the fatigue-induced decline in pH_i_ was less marked in *Tfam* KO than in WT mice (Figure 5, E-F). None of the measured metabolic changes during fatiguing stimulation showed any difference between CsA-treated and untreated *Tfam* KO mice.

**Figure 5.**
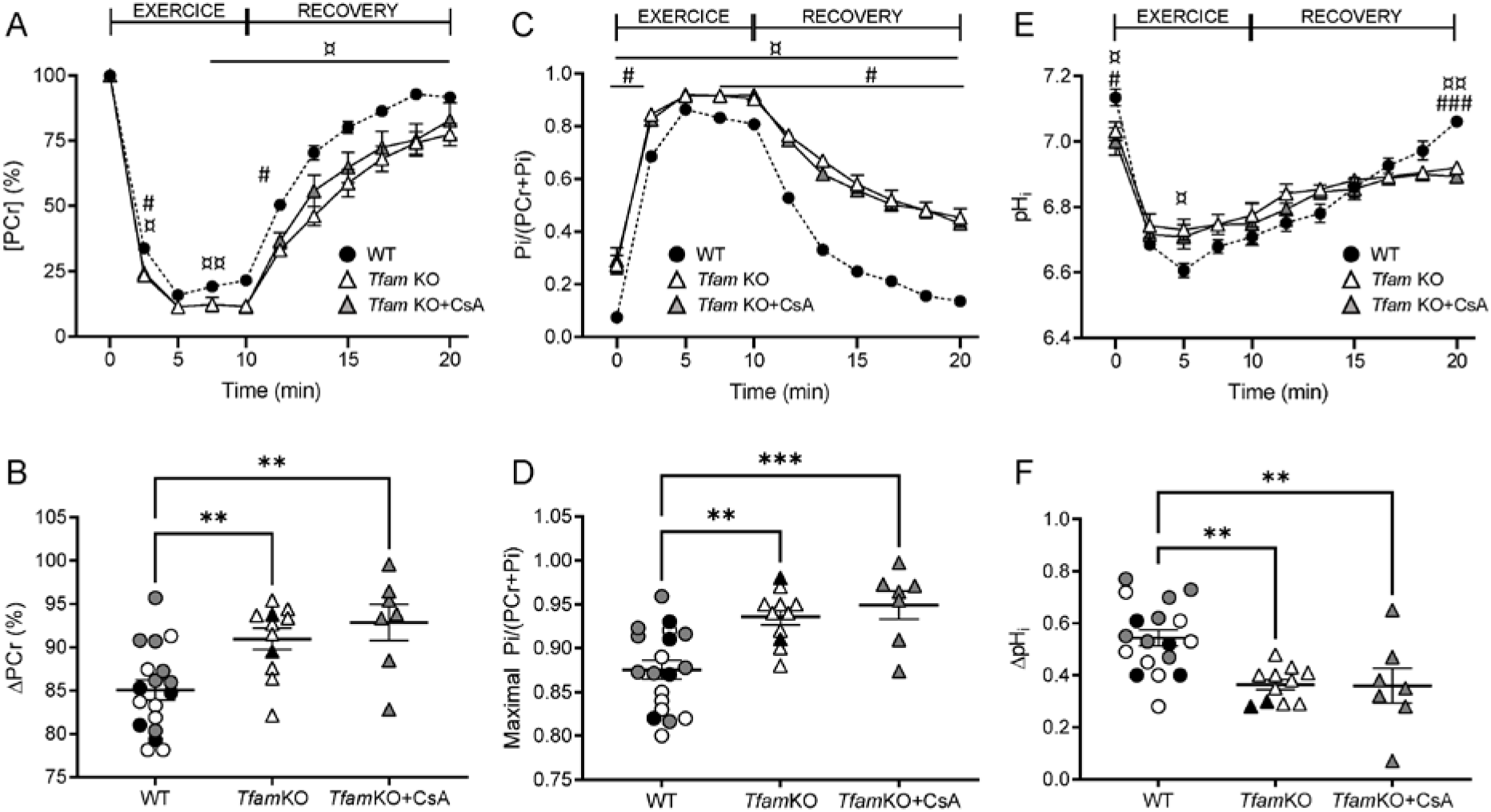
Administration of CsA for 4 weeks had no effect on muscle energetics during exercise in *Tfam* KO mice. **(A)** Phosphocreatine levels throughout the stimulation period and during the recovery after the stimulation. **(B)** Magnitude of phosphocreatine reduction during the stimulation period. **(C)** Inorganic phosphate levels during fatiguing stimulation and recovery. **(D)** Maximal inorganic phosphate reached during fatiguing protocol. **(E)** pH_i_ during stimulation and recovery periods. **(F)** Magnitude of pH decrease during the stimulation period. Values are mean±SEM for panels A, C and E. Data presented as individual values and mean±SEM for other panels. WT, n=19; *Tfam* KO, n=11; *Tfam* KO+CsA, n=10. White=mouse without pump; Black=mouse with pump+placebo; Grey=mouse with pump+CsA, for panels B, D and F. Significant difference between WT and *Tfam* KO ^¤^ and ^¤¤^ *P*<0.05 and 0.01, respectively. Significant difference between WT and *Tfam* KO+CsA ^#^ and ^###^ *P*<0.05 and 0.01, respectively. Significant difference ** and *** *P*<0.01 and 0.001, respectively. Two-way ANOVA with repeated measures on time and Sidak’s post hoc test for time-course of phosphocreatine, inorganic phosphate and pH_i_ and One-way ANOVA and Tuckey’s post hoc test for all other comparisons. PCr: phosphocreatine; Pi: inorganic phosphate; pH_i_: intracellular pH.

^31^P-MRS measurements were continued for 10 min of recovery after the fatiguing stimulation. The rate and total extent of PCr recovery were lower in *Tfam* KO than in WT mice (Figure 6, A-C). CsA treatment of *Tfam* KO mice did not result in any marked difference in the measured PCr recovery parameters as compared to untreated *Tfam* KO mice. Nevertheless, there was a trend towards improvement considering that these measurements were only statistically significant different between untreated *Tfam* KO and WT mice. At the end of the 10 min recovery period, Pi and pH_i_ had returned to their pre-fatigue resting values with higher Pi levels and lower pH_i_ in *Tfam* KO than in WT mice (Figure 6, D-E). Taken together, our results show no obvious beneficial effects of CsA treatment on the defective mitochondrial energy production in *Tfam* KO mice.

**Figure 6.**
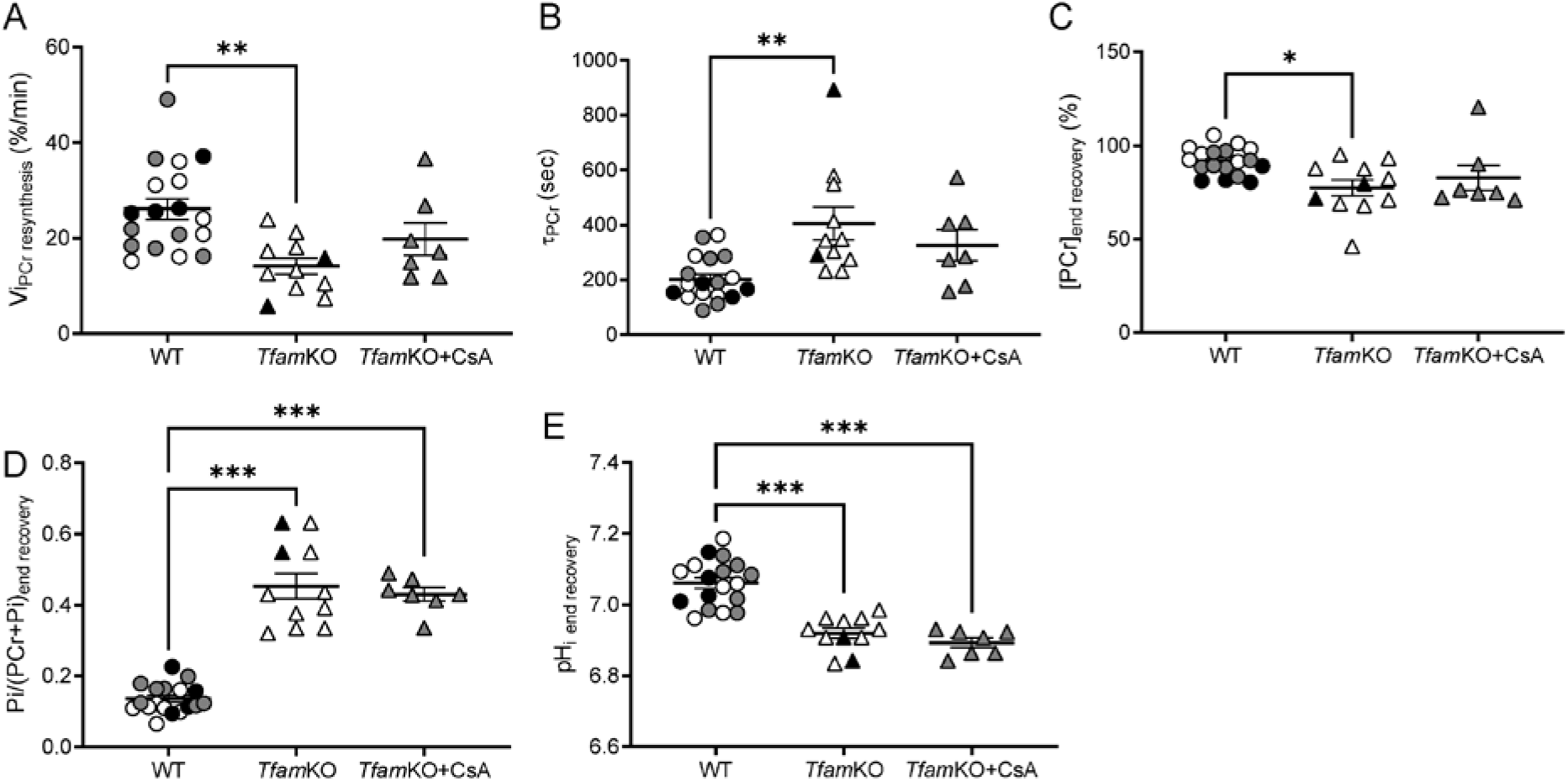
Mitochondrial oxidative capacity was unchanged in *Tfam* KO mice following 4-week administration of CsA. Phosphocreatine **(A)** and inorganic phosphate **(B)** at the end of the 10-min recovery period. **(C)** Initial rate of phosphocreatine resynthesis after the stimulation bout. **(D)** Time constant of phosphocreatine recovery during the 10 min recovery phase after exercise. **(E)** pH_i_ at the end of the 10-min recovery period. Data presented as individual values and mean±SEM. WT, n=19; *Tfam* KO, n=11; *Tfam* KO+CsA, n=10. White=mouse without pump; Black=mouse with pump+placebo; Grey=mouse with pump+CsA. Significant difference *, ** and *** *P*<0.05, 0.01 and 0.001, respectively. One-way ANOVA and Tuckey’s post hoc test for panels A and E; Kruskal-Wallis test and Dunn’s post hoc test for all other comparisons. PCr: phosphocreatine; Pi: inorganic phosphate; pH_i_: intracellular pH; Vi_PCr resynthesis_: initial rate of phosphocreatine resynthesis; τ_PCr_: time constant of phosphocreatine recovery.

## Discussion

In the present study, physiological experiments were performed *in vivo* in *Tfam* KO mice, *i. e*. a mouse model of severe mitochondrial myopathy. Our findings confirm and extend previous results showing that CsA treatment can delay the disease progression into the terminal phase which is characterized by a rapidly developing muscle weakness, weight loss and ultimately premature death (15). Unexpectedly, this beneficial effect of CsA treatment occurred despite no obvious improvement of the compromised mitochondrial energy production (11).

The effects of 4-week CsA administration in *Tfam* KO mice started at ~12 weeks, an age where muscles of *Tfam* KO mice had not yet entered the terminal stage with rapidly progressing muscle atrophy and weakness. The disease process was faster in untreated *Tfam* KO mice as demonstrated, for instance, by the untreated mice reaching their maximum body weight 3 weeks earlier than the CsA-treated mice. Nevertheless, there was no difference in hindlimb muscle volume or tetanic force production in the unfatigued state between CsA-treated and untreated *Tfam* KO mice at the ages of the present *in vivo* testing. This is in sharp contrast with previous results obtained when both untreated and CsA-treated *Tfam* KO mice were tested at the same age of 16 weeks. At this age, lean hindlimb mass and tetanic force were markedly lower in the untreated group (15). However, in this study untreated *Tfam* KO mice could not be tested at the same age than the CsA-treated *Tfam* KO mice. The untreated *Tfam* KO mice started to lose weight after the intervention period started. Knowing that terminal phase occurs between 12 and 16 weeks and given that we previously reported progression of muscle atrophy and force production between 11 and 14 weeks (11, 12), we could expect worsening of these parameters after 14 weeks. For this reason, experiments were performed before untreated *Tfam* KO mice reached ethical endpoint (*i. e*. >15% loss of body weight). Therefore, *in vivo* investigations were performed earlier (*i. e*. at an age of ~14 weeks) in the untreated *Tfam* KO group, while the CsA-treated mice were tested at the end of this intervention period, *i. e*. when mice were ~16 weeks old. Our *in vivo* investigations, which require anesthesia, would have certainly not been feasible on older untreated *Tfam* KO mice due to muscle weakness. In the untreated *Tfam* KO mice muscle weakness is not yet installed, probably because of the younger age as compared to previous results (15). Muscle weakness was also absent in the CsA-treated *Tfam* KO mice despite the older age which suggests that CsA may help to prevent loss in force production. In addition, the absence of worsening muscle atrophy in CsA-treated *Tfam* KO mice likely indicates a protective effect of CsA on muscle in this study.

*In vivo* comparative analysis of younger untreated than CsA-treated *Tfam* KO mice allowed us to assess muscle fatigue and the accompanying metabolic changes at comparable starting forces. On that basis, one could avoid a bias effect related to a markedly lower initial force which could also be linked to a decreased energy demand during fatigue induction in the untreated group. This prevented a markedly lower starting force, and hence decreased energy demand during induction of fatigue in the untreated group (13). Force decline was markedly faster during the first part of fatiguing stimulation in *Tfam* KO than in WT mice and this accelerated fatigue development in *Tfam* KO mice was not affected by CsA treatment. Towards the end of fatiguing stimulation, force reached a relatively stable level at ~20% of the original force in both *Tfam* KO and WT mice, which is likely due to force being maintained at this low level by slow-twitch type 1 muscle fibers that are not knocked out in the present fast-twitch fiber-selective *Tfam* KO model (12).

Our ^31^P-MRS measurements revealed differences in metabolites content between *Tfam* KO and WT muscles at rest (lower [PCr]/[ATP] ratio, higher [Pi] and lower pH_i_), thereby illustrating severe mitochondrial deficiency in *Tfam* KO muscles. The mitochondrial oxidative capacity of muscle fibers is of key importance in order to limit force decline during the present type of fatiguing stimulation with repeated contractions elicited by electrical muscle stimulation (19, 20). The fatigue-induced metabolic changes (decreased [PCr] accompanied by the [Pi] accumulation and the intracellular acidosis) were generally larger and recovered more slowly in *Tfam* KO than in WT mice. Hence, this indicates a lower mitochondrial oxidative capacity and larger dependency on anaerobic metabolism in *Tfam* KO muscle. The only exception from this was a smaller decline in pH_i_ in *Tfam* KO than in WT muscles during induction of fatigue, which might relate to the larger breakdown of PCr in *Tfam* KO muscle resulting in increased uptake of H^+^ via the creatine kinase reaction (PCr + ADP + H^+^ → Cr + ATP → Pi + Cr + ADP) (21). A partial resistance to intracellular acidosis has been observed in exercising muscle of patients with mitochondrial diseases and suggested to be an adaptation of intracellular buffering systems in response to chronic overproduction of lactate (22, 23). Importantly, CsA treatment did not limit the metabolic deficiency in *Tfam* KO muscle either at rest or during fatigue, which implies no beneficial effect of CsA treatment on mitochondrial energy production and fits with no CsA-mediated improvement of force production during fatigue.

Treatment with CsA as well as other cyclophilin inhibitors or genetic ablation of CypD have been related to marked positive effects in several mouse disease models where defective mitochondrial function was a key feature (15, 24–28). In addition, positive effects have also been reported in patients with CypD-dependent mitochondrial defects due to collagen VI mutations (29). Moreover, the protein expression of CypD has been shown to be increased in patients with mitochondrial DNA depletion or mutations (15), in a recently described mouse model with impaired mitochondrial function due to skeletal muscle-specific deletion of nicotinamide phosphoribosyltransferase (NAMPT) resulting in severely decreased NAD^+^ and NADH levels (24), and in the present *Tfam* KO model (15). Here we show that in *Tfam* KO mice, CsA treatment delays the disease progression into the terminal phase with severe muscle weakness without having any obvious beneficial effects on mitochondrial respiratory function. Our results are consistent with the results from the NAMPT-deficient mouse model where CsA treatment reduced muscle fiber damage and improved survival without affecting the energetic stress (24).

The main limitation of this study is the age difference between the untreated and CsA-treated *Tfam* KO mice. Indeed, as mentioned above, untreated *Tfam* KO mice were tested at an earlier age than the CsA-treated *Tfam* KO for ethical and practical reasons. As the disease progress towards terminal phase from 12 weeks (rapid muscle atrophy and weakness), the phenotype would have definitely been worse at 16 weeks in the untreated *Tfam* KO mice than at the actual 14-week of age. Considering that our results show that CsA delays disease progression in *Tfam* KO mice, the effects of CsA might be played down. Indeed, if the two *Tfam* KO groups were tested at the same age, positive effects of CsA on muscle and/or mitochondrial function of *Tfam* KO mice would have probably been observed. In addition, in *Tfam* KO mice, energy production defects start earlier than muscle weakness and muscle atrophy, and we previously showed that they were already apparent at 11 weeks (11). Therefore, it cannot be excluded that CsA may have had protective effects on mitochondrial function, and therefore on muscle fatigue, if treatment was initiated before severe energy defects were established.

In conclusion, CypD inhibitors have the potential to counteract the devastating muscle weakness in patients with mitochondrial myopathies. This effect likely occurs by preventing deleterious effects triggered by excessive mitochondrial Ca^2+^ uptake rather than by improving mitochondrial energy production.

## Materials and Methods

### Animals, groups and treatment

*Tfam* KO mice and control littermates (referred to as wild type, WT) were used for experiments. The fast-twitch skeletal muscle-specific *Tfam* KO mouse model was generated as previously described (12). *Tfam* KO mice and WT mice were randomly assigned to either the treated or untreated group. Treated mice received CsA (120 μg/day) for 4 weeks via osmotic pumps implanted subcutaneously on the back under anesthesia (2.5% isoflurane in 33% O_2_ and 66% NO_2_). CsA treatment of *Tfam* KO mice started at 12-13 weeks old. None of the CsA-treated *Tfam* KO mice had lost weight before pump implantation. Untreated mice either had implanted osmotic pumps with vehicle (*i. e*. containing only saline: 2 of 15 *Tfam* KO mice; 4 of 11 WT mice) or were left without any treatment. We did not observe any adverse effects of the pump implantation either in WT mice or in *Tfam* KO mice; thus, data have been pooled but different colors were used for individual data points from mice with (grey) and without (white) pumps implanted.

The duration of the CsA treatment spanned from 12 to 16 weeks of age, covering the period when the myopathy goes from mild to terminal. Experiments on CsA-treated mice were performed at the end of the treatment period, *i. e*. 16 weeks. Untreated *Tfam* KO mice would frequently not survive to 16 weeks of age. Experiments on untreated *Tfam* KO mice were therefore performed when they started to lose weight indicating progression of the disease towards the end stage (20% loss in maximum body weight).

### Experimental design

Body weight was measured every 2 to 3 days until mice were subjected to a strictly noninvasive *in vivo* anatomical and functional measurements in muscles of the left hindlimb. These *in vivo* experiments were performed after 4 weeks of intervention or earlier if mice showed an accelerated decline in body weight, *i. e*. a sign of approaching the terminal disease state. The contractile performance of plantar flexor muscles [*gastrocnemius* (Gas), *soleus* (Sol), and *plantaris*] was assessed by measuring the maximal tetanic force in the unfatigued state and force production during a fatiguing stimulation (see below). Metabolic changes were evaluated before, during and after fatiguing stimulation using phosphorus-31 magnetic resonance spectroscopy (^31^P-MRS). Before the fatiguing protocol, the volume of hindlimb muscles was quantified by magnetic resonance imaging (MRI). Mice were euthanized by cervical dislocation at the end of the *in vivo* force and magnetic resonance experiments.

### Magnetic resonance and force output measurements

Mice were anesthetized (4% isoflurane in 33% O_2_ (0.5 l/min) and 66% NO_2_ (1 l/min)) and placed supine in a cradle designed in our laboratory for the strictly non-invasive functional investigation of the left hindlimb muscles. A custom-built facemask was incorporated into the cradle and was used to maintain a prolonged anesthesia throughout the experiment (1.75% isoflurane in 33% O_2_ (0.2 l/min) and 66% NO_2_ (0.4 l/min)). The hindlimb was centered inside a ^1^H imaging coil and the belly of the Gas muscle was located above a ^31^P-MRS surface coil. The foot was positioned on the pedal of a home-made ergometer with a 90° flexion ankle joint coupled to a force transducer. The analog electrical signal coming out from the force transducer was amplified with a home-built amplifier (Operational amplifier AD620; Analog Devices, Norwood, MA, USA), converted to a digital signal, monitored and recorded on a personal computer using the Powerlab 35/series system (AD Instruments, Oxford, United Kingdom). Two rod-shaped surface electrodes integrated into the cradle and connected to an electrical stimulator (Digitimer, Ltd., Model DS7A, Welwyn Springs, UK) were placed on the left hindlimb, one at the heel level and the other one just above the knee joint. Skeletal muscle contractions were achieved by non-invasive transcutaneous electrical stimulation elicited with square-wave pulses (0.5 ms duration). The electrical current required to obtain maximal muscle activation was in each mouse determined by progressively increasing the pulse intensity until there was no further increase in peak twitch force. Maximal tetanic force of the plantar flexor muscles was assessed in response to a 150 Hz tetanic stimulation train with a duration of 0.75 s. Force production of the plantar flexor muscles was also measured during a fatigue protocol (80 contractions at 40 Hz, 1.5 s on - 6 s off) (11, 30). In order to assess contractile function independent of muscle size, force values have been scaled to the corresponding muscle volume values.

MR experiments were performed using a 47/30 Biospec Avance MR system (Bruker, Karlsruhe, Germany) equipped with a 120-mm BGA12SL (200 mT/m) gradient insert. Anatomical MRI data were acquired at rest, *i.e*. before functional measurements. Fifteen consecutive contiguous axial slices (thickness=0.7 mm), covering the region from the knee to the ankle, were selected across the lower hindlimb. Rapid acquisition with relaxation enhancement (RARE) images (RARE factor=4, effective echo time=22.5 ms, actual echo time=10.6 ms, repetition time=1000 ms, number of averages=10, number of repetitions=1, field of view=4.2×4.2 cm, matrix size=256×192, and acquisition time=8 min 00 s) were recorded. ^31^P spectra (8-kHz sweep width; 2048 data points) from the posterior hindlimb muscles region were acquired continuously throughout the standardized rest-fatigue-recovery protocol. A total of 800 partially saturated (repetition time=2 s) free induction decays (FID) were recorded.

### Data processing

#### Anatomical imaging

Images were analyzed using FSLview (FMRIB, Oxford, UK) (31). The border of the whole hindlimb muscle area was manually delineated in the two slices located on the proximal and distal parts. Segmentations of the missing intermediate slices was automatically performed on the basis of registration procedures as described previously (32). The volume of the hindlimb muscles (mm^3^) was calculated as the sum of the volume of the ten consecutive largest slices.

#### Contractile performance

The peak force of each contraction was measured using the LabChart software (AD Instruments, Oxford, United Kingdom). For the fatigue protocol, the tetanic force was averaged every 5 contractions and the total force production was computed as the sum of each individual peak forces measured during the fatiguing stimulation. The C_50_ corresponds to the contraction number at which the force amplitude was half of the maximal force recorded during the fatigue protocol. Force was normalized with respect to the hindlimb muscles volume to obtain the specific force (in mN/mm^3^).

#### Metabolism

^31^P-MRS data were processed using a proprietary software developed using IDL (Interactive Data Language, Research System, Inc., Boulder, CO, USA) (33). The first 180 FID were acquired at rest and summed together (n=1, time resolution=6 min). The next 320 FID were acquired during the stimulation period and summed by blocks of 80 during the stimulation procedure (n=4, time resolution=160 s). The last 300 FID were acquired during the recovery period and summed by blocks of 50 (n=6, time resolution=100 s). The relative concentrations of high-energy phosphate metabolites (phosphocreatine (PCr) and inorganic phosphate (Pi)) were obtained by a time-domain fitting routine using the AMARES-MRUI Fortran code and appropriate prior knowledge of the ATP multiplets. In order to correct for potential movement artifacts during fatiguing stimulation, Pi values were normalized to the sum of PCr and Pi values, which were considered as constant (34). Intracellular pH (pH_i_) was calculated from the chemical shift of the Pi signal relative to PCr (35). The resting phosphocreatine to ATP ratio was calculated from the peak areas of the phosphocreatine and β-ATP of the spectrum acquired at rest. Changes during the fatiguing protocol (ΔPCr, maximal Pi/(Pi+PCr) and ΔpH_i_) and recovery after exercise (initial rate of PCr resynthesis (Vi _PCr resynthesis_) and PCr recovery time constant (τ_PCr_)) were extracted from the time-courses of PCr, Pi and pH_i_ (11).

### Statistical analysis

Data are presented as mean ± SEM. Statistical analyses were performed using GraphPad Prism software version 9.1.2. Normality was checked using a Shapiro-Wilk test. Parametric tests were performed when data were normally distributed. If data were not normally distributed, non-parametric tests were used. One-way ANOVA was applied for comparisons between WT, *Tfam* KO and *Tfam* KO+CsA. Two-way ANOVA with repeated measures on time or contraction number was used to assess force production and metabolic changes during induction of fatigue and the subsequent recovery. When a main effect or a significant interaction was found, Tukey post-hoc analysis or Sidak post-hoc was performed. Student’s unpaired *t*-test or Mann-Whitney test was used for all other comparisons. Statistical significance was accepted when *P<*0.05.

### Study approval

All experiments were conducted in agreement with the French guidelines for animal care and in accordance with the European Convention for the Protection of Vertebrate Animals used for Experimental and other Scientific Purposes, and institutional guidelines n° 86/609/CEE November 24, 1986. All animal experiments were approved by the Institutional Animal Care Committee of Aix-Marseille University (permit number #12522-2017121119249655 v1). Mice were housed in an environment-controlled facility (12–12 h light-dark cycle at 22 °C) and received water and standard food *ad libitum*.

## Author contributions

CG conceived the project. BC, CG and EP performed the experiments. BC, IV, AO, CG analyzed the data. CG, BC and HW wrote the manuscript. MB, JG, DB provided support and edited the manuscript. CG and DB supervised the work.

## Acknowledgements

This work was funded by the AFM-Téléthon (trampoline grant #19963 to CG). CRMBM is member of France Life Imaging network (grant ANR-11-INBS-0006).

## Notes

**Conflict-of-interest statement:** The authors have declared that no conflict of interest exists.

### Competing Interest Statement

The authors have declared no competing interest.

## References

1. Thorburn DR. Mitochondrial disorders: prevalence, myths and advances. J Inherit Metab Dis. 2004;27(3):349–362.

2. Petty RK, Harding AE, and Morgan-Hughes JA. The clinical features of mitochondrial myopathy. Brain. 1986;109 (Pt 5):915–938.

3. Taivassalo T, Jensen TD, Kennaway N, DiMauro S, Vissing J, and Haller RG. The spectrum of exercise tolerance in mitochondrial myopathies: a study of 40 patients. Brain. 2003;126(Pt 2):413–423.

4. Ahmed ST, Craven L, Russell OM, Turnbull DM, and Vincent AE. Diagnosis and Treatment of Mitochondrial Myopathies. Neurotherapeutics. 2018;15(4):943–953.

5. Taivassalo T, and Haller RG. Exercise and training in mitochondrial myopathies. Med Sci Sports Exerc. 2005;37(12):2094–2101.

6. Ahola-Erkkila S, Carroll CJ, Peltola-Mjosund K, Tulkki V, Mattila I, Seppanen-Laakso T, Oresic M, Tyynismaa H, and Suomalainen A. Ketogenic diet slows down mitochondrial myopathy progression in mice. Hum Mol Genet. 2010;19(10):1974–1984.

7. Bendahan D, Desnuelle C, Vanuxem D, Confort-Gouny S, Figarella-Branger D, Pellissier JF, Kozak-Ribbens G, Pouget J, Serratrice G, and Cozzone PJ. 31P NMR spectroscopy and ergometer exercise test as evidence for muscle oxidative performance improvement with coenzyme Q in mitochondrial myopathies. Neurology. 1992;42(6):1203–1208.

8. Al Jasmi F, Al Zaabi N, Al-Thihli K, Al Teneiji AM, Hertecant J, and El-Hattab AW. Endothelial Dysfunction and the Effect of Arginine and Citrulline Supplementation in Children and Adolescents With Mitochondrial Diseases. J Cent Nerv Syst Dis. 2020;12:1179573520909377.

9. Pfeffer G, Horvath R, Klopstock T, Mootha VK, Suomalainen A, Koene S, Hirano M, Zeviani M, Bindoff LA, Yu-Wai-Man P, et al. New treatments for mitochondrial disease-no time to drop our standards. Nat Rev Neurol. 2013;9(8):474–481.

10. Zweers H, van Wegberg Amj, Janssen MCH, and Wortmann SB. Ketogenic diet for mitochondrial disease: a systematic review on efficacy and safety. Orphanet J Rare Dis. 2021;16(1):295.

11. Chatel B, Ducreux S, Harhous Z, Bendridi N, Varlet I, Ogier AC, Bernard M, Gondin J, Rieusset J, Westerblad H, et al. Impaired aerobic capacity and premature fatigue preceding muscle weakness in the skeletal muscle Tfam KO mouse model. Dis Model Mech. 2021;14(9):dmm048981.

12. Wredenberg A, Wibom R, Wilhelmsson H, Graff C, Wiener HH, Burden SJ, Oldfors A, Westerblad H, and Larsson NG. Increased mitochondrial mass in mitochondrial myopathy mice. Proc Natl Acad Sci U S A. 2002;99(23):15066–15071.

13. Aydin J, Andersson DC, Hanninen SL, Wredenberg A, Tavi P, Park CB, Larsson NG, Bruton JD, and Westerblad H. Increased mitochondrial Ca2+ and decreased sarcoplasmic reticulum Ca2+ in mitochondrial myopathy. Hum Mol Genet. 2009;18(2):278–288.

14. Briston T, Selwood DL, Szabadkai G, and Duchen MR. Mitochondrial Permeability Transition: A Molecular Lesion with Multiple Drug Targets. Trends Pharmacol Sci. 2019;40(1):50–70.

15. Gineste C, Hernandez A, Ivarsson N, Cheng AJ, Naess K, Wibom R, Lesko N, Bruhn H, Wedell A, Freyer C, et al. Cyclophilin D, a target for counteracting skeletal muscle dysfunction in mitochondrial myopathy. Hum Mol Genet. 2015;24(23):6580–6587.

16. Yamada T, Ivarsson N, Hernandez A, Fahlstrom A, Cheng AJ, Zhang SJ, Bruton JD, Ulfhake B, and Westerblad H. Impaired mitochondrial respiration and decreased fatigue resistance followed by severe muscle weakness in skeletal muscle of mitochondrial DNA mutator mice. J Physiol. 2012;590(23):6187–6197.

17. Finsterer J. Update Review about Metabolic Myopathies. Life (Basel). 2020;10(4):43.

18. Tarnopolsky MA. Metabolic Myopathies. Continuum (Minneap Minn). 2016;22(6, Muscle and Neuromuscular Junction Disorders):1829–1851.

19. Westerblad H, Bruton JD, and Katz A. Skeletal muscle: energy metabolism, fiber types, fatigue and adaptability. Exp Cell Res. 2010;316(18):3093–3099.

20. Allen DG, Lamb GD, and Westerblad H. Skeletal muscle fatigue: cellular mechanisms. Physiol Rev. 2008;88(287-332.

21. Hultman E, and Sahlin K. Acid-base balance during exercise. Exerc Sport Sci Rev. 1980;8:41–128.

22. Argov Z, Bank WJ, Maris J, Peterson P, and Chance B. Bioenergetic heterogeneity of human mitochondrial myopathies: phosphorus magnetic resonance spectroscopy study. Neurology. 1987;37(2):257–262.

23. Arnold DL, Taylor DJ, and Radda GK. Investigation of human mitochondrial myopathies by phosphorus magnetic resonance spectroscopy. Ann Neurol. 1985;18(2):189–196.

24. Basse AL, Agerholm M, Farup J, Dalbram E, Nielsen J, Ortenblad N, Altintas A, Ehrlich AM, Krag T, Bruzzone S, et al. Nampt controls skeletal muscle development by maintaining Ca(2+) homeostasis and mitochondrial integrity. Mol Metab. 2021;53:101271.

25. Palma E, Tiepolo T, Angelin A, Sabatelli P, Maraldi NM, Basso E, Forte MA, Bernardi P, and Bonaldo P. Genetic ablation of cyclophilin D rescues mitochondrial defects and prevents muscle apoptosis in collagen VI myopathic mice. Hum Mol Genet. 2009;18(11):2024–2031.

26. Zulian A, Rizzo E, Schiavone M, Palma E, Tagliavini F, Blaauw B, Merlini L, Maraldi NM, Sabatelli P, Braghetta P, et al. NIM811, a cyclophilin inhibitor without immunosuppressive activity, is beneficial in collagen VI congenital muscular dystrophy models. Hum Mol Genet. 2014;23(20):5353–5363.

27. Irwin WA, Bergamin N, Sabatelli P, Reggiani C, Megighian A, Merlini L, Braghetta P, Columbaro M, Volpin D, Bressan GM, et al. Mitochondrial dysfunction and apoptosis in myopathic mice with collagen VI deficiency. Nat Genet. 2003;35(4):367–371.

28. Millay DP, Sargent MA, Osinska H, Baines CP, Barton ER, Vuagniaux G, Sweeney HL, Robbins J, and Molkentin JD. Genetic and pharmacologic inhibition of mitochondrial-dependent necrosis attenuates muscular dystrophy. Nat Med. 2008;14(4):442–447.

29. Merlini L, Angelin A, Tiepolo T, Braghetta P, Sabatelli P, Zamparelli A, Ferlini A, Maraldi NM, Bonaldo P, and Bernardi P. Cyclosporin A corrects mitochondrial dysfunction and muscle apoptosis in patients with collagen VI myopathies. Proc Natl Acad Sci U S A. 2008;105(13):5225–5229.

30. Gineste C, Ogier AC, Varlet I, Hourani Z, Bernard M, Granzier H, Bendahan D, and Gondin J. In vivo characterization of skeletal muscle function in nebulin-deficient mice. Muscle Nerve. 2020;61(3):416–424.

31. Jenkinson M, Beckmann CF, Behrens TE, Woolrich MW, and Smith SM. Fsl. Neuroimage. 2012;62(2):782–790.

32. Ogier A, Sdika M, Foure A, Le Troter A, and Bendahan D. Individual muscle segmentation in MR images: A 3D propagation through 2D non-linear registration approaches. Annu Int Conf IEEE Eng Med Biol Soc. 2017;2017:317–320.

33. Le Fur Y, Nicoli F, Guye M, Confort-Gouny S, Cozzone PJ, and Kober F. Grid-free interactive and automated data processing for MR chemical shift imaging data. Magma. 2010;23(1):23–30.

34. Chance B, Eleff S, Leigh JS, Jr., Sokolow D, and Sapega A. Mitochondrial regulation of phosphocreatine/inorganic phosphate ratios in exercising human muscle: a gated 31P NMR study. Proc Natl Acad Sci U S A. 1981;78(11):6714–6718.

35. Moon RB, and Richards JH. Determination of intracellular pH by 31P magnetic resonance. J Biol Chem. 1973;248(20):7276–7278.

