## Supplementary Data for "Cyclosporin A delays the terminal disease stage in *Tfam* KO mice without improving mitochondrial energy production"

**Supplementary Table 1.** Summary of the anatomic, metabolic, and force measurements in mice that did not undergo surgery and mice that underwent surgery and received placebo

| | WT | | *Tfam* KO | |
|---|---|---|---|---|
| | **-surgery** | **+surgery** | **-surgery** | **+surgery** |
| **Hindlimb muscles volume ($mm^3$)** | 200±7 | 186±11 | 173±9 | 164±15 |
| **Force – 150 Hz ($mN/mm^3$)** | 1.88±0.12 | 1.83±0.13 | 1.77±0.06 | 1.94±0.16 |
| **Total force – exercise ($mN/mm^3$)** | 10.3±0.4 | 9.0±0.8 | 7.3±0.6 | 6.9±2.3 |
| **$C_{50}$ (number of contractions)** | 28±3 | 23±3 | 14±1 | 15±3 |
| **$PCr/ATP_{rest}$** | 2.41±0.10 | 1.90±0.22 | 1.96±0.10 | 2.30±0.07 |
| **$Pi/(PCr+Pi)_{rest}$** | 0.089±0.016 | 0.065±0.011 | 0.25±0.04 | 0.39±0.10 |
| **$pH_{i\ rest}$** | 7.10±0.03 | 6.99±0.04 | 7.06±0.03 | 6.92±0.03 |
| **$Pi/(PCr+Pi)_{max}$** | 0.85±0.02 | 0.88±0.02 | 0.93±0.01 | 0.95±0.04 |
| **$Vi_{PCr\ degradation}$ (%/min)** | 56.4±6.5 | 56.5±7.8 | 91.2±16.0 | 89.1±11.6 |
| **ΔPCr** | 83±2 | 83±1 | 91±2 | 92±2 |
| **$\Delta pH_i$** | 0.50±0.05 | 0.48±0.05 | 0.38±0.02 | 0.29±0.01 |
| **$Vi_{PCr\ resynthesis}$ (%/min)** | 25.0±3.1 | 28.5±2.9 | 14.9.0±1.9 | 10.8±5.1 |
| **$\tau_{PCr}$ (sec)** | 211±32 | 161±11 | 364±43 | 592±300 |

No surgery: WT-surgery, n=7; TfamKO-surgery, n=11. Surgery: WT+surgery, n=4; TfamKO+surgery, n=2. No significant difference was found between the two groups of WT mice and the two groups of *Tfam* KO mice for any of the parameters measured. Values are mean±SEM. $C_{50}$: number of contractions from which force production was 50% of the maximal force; PCr: phosphocreatine; ATP: adenosine triphosphate; $Vi_{PCr\ degradation}$: initial rate of PCr degradation; $Vi_{PCr\ resynthesis}$: initial rate of PCr resynthesis; $\tau_{PCr}$: time constant of PCr recovery;

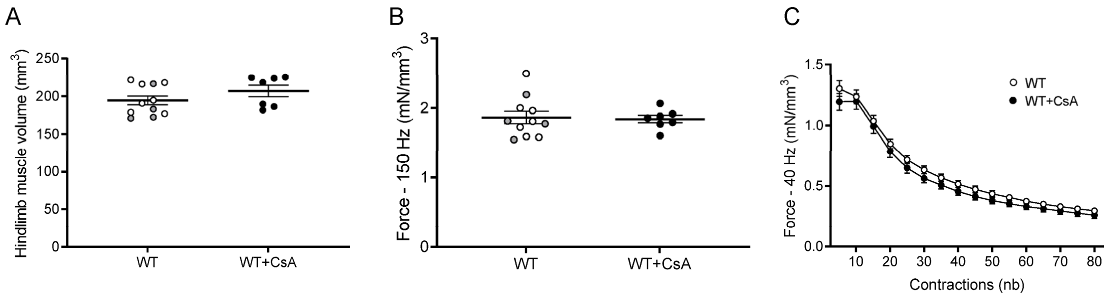


**Supplementary Figure 1** Muscle anatomy and fatigue were unchanged in WT mice following CsA administration. **(A)** Volume of hindlimb muscles. **(B)** Maximal in vivo specific force. **(C)** *In vivo* specific force during 80-stimulation exercise. WT, n=11; WT+CsA: n=8. White=mouse without pump; Black=mouse with pump+placebo; Grey=mouse with pump+CsA for panels A and B. Data presented as individual values and mean±SEM in panels A and B. Values are mean±SEM for panel C. No significant difference was found for any measurements. Unpaired t-test was applied for panels A & B and two-way ANOVA with repeated measures on time or contraction number and Sidak's post-hoc test were used for panel C.

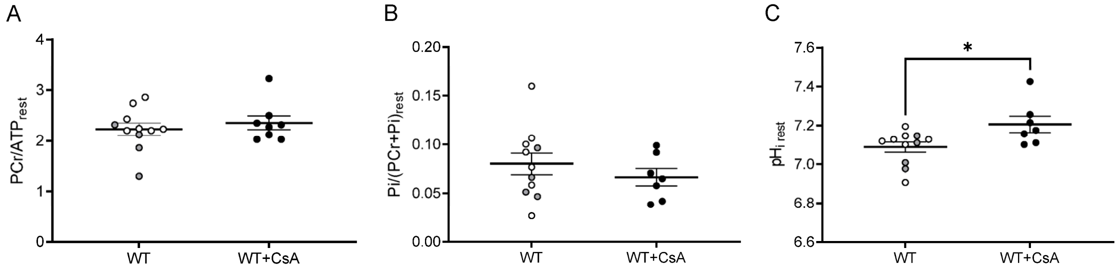


**Supplementary Figure 2** Muscle metabolism at rest was similar in WT mice with and without CsA. Phosphocreatine to ATP ratio **(A)**, inorganic phosphate **(B)** and $pH_i$ **(C)** measured at rest. WT, n=11; WT+CsA: n=8. White=mouse without pump; Black=mouse with pump+placebo; Grey=mouse with pump+CsA. Data presented as individual values and mean±SEM. No significant difference between groups; Mann-Whitney test for panel A; unpaired t-test for panels B and C. PCr: phosphocreatine; ATP: Adenosine tri-phosphate; Pi: inorganic phosphate; $pH_i$: intracellular pH.

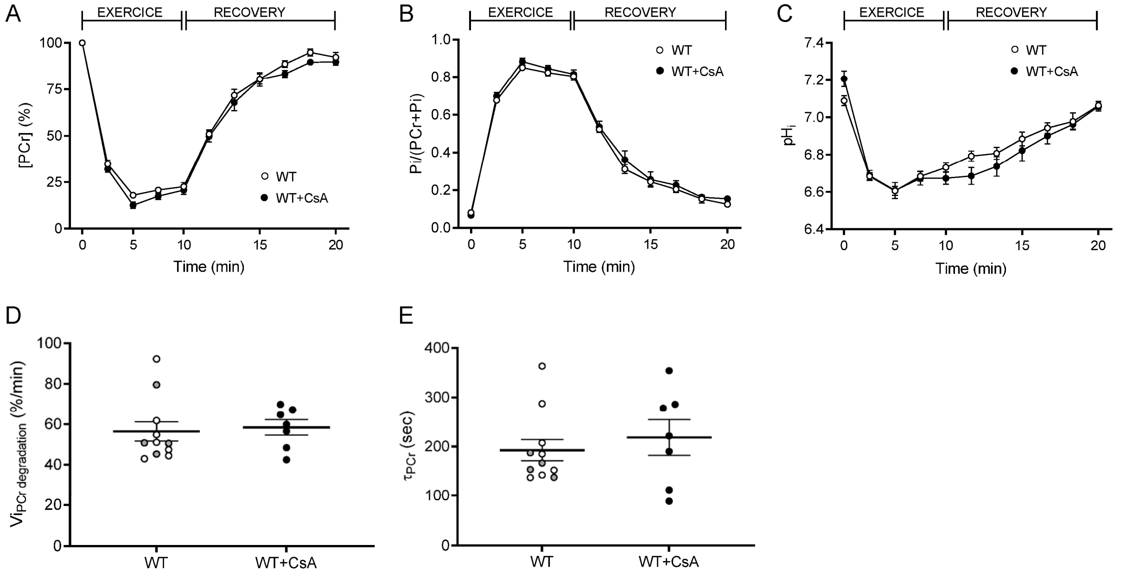


**Supplementary Figure 3** Muscle metabolism during exercise and recovery was similar in WT mice with and without CsA. Phosphocreatine **(A)**, inorganic phosphate **(B)** and pH$_i$ **(C)** levels throughout the stimulation period and during the recovery after the stimulation. **(D)** Initial rate of phosphocreatine degradation. **(E)** Time constant of phosphocreatine recovery during the 10 min recovery phase after exercise. WT, n=11; WT+CsA: n=8. White=mouse without pump; Black=mouse with pump+placebo; Grey=mouse with pump+CsA for panels D and E. Values are mean±SEM in panels A, B & C. Data presented as individual values and mean±SEM in panels D and E. No significant difference between groups; two-way ANOVA with repeated measures on time and Sidak's post-hoc test for panels A, B & C; Mann-Whitney test for panels D and E. PCr: phosphocreatine; Pi: inorganic phosphate; pH$_i$: intracellular pH; Vi $_{PCr\ degradation}$: initial rate of phosphocreatine breakdown; $\tau_{PCr}$: time constant of phosphocreatine recovery.
